# Neural signatures of automatic repetition detection in temporally regular and jittered acoustic sequences

**DOI:** 10.1101/2023.04.11.536415

**Authors:** Hanna Ringer, Erich Schröger, Sabine Grimm

## Abstract

Detection of repeating patterns within continuous sound streams is crucial for efficient auditory perception. Previous studies demonstrated a remarkable sensitivity of the human auditory system to periodic repetitions in randomly generated sounds. Automatic repetition detection was reflected in different EEG markers, including sustained activity, neural synchronisation, and event-related responses to pattern occurrences. The current study investigated how listeners’ attention and the temporal regularity of a sound modulate repetition perception, and how this influence is reflected in different EEG markers that were previously suggested to subserve dissociable functions. We reanalysed data of a previous study in which listeners were presented with random acoustic sequences with and without repetitions of a certain sound segment. Repeating patterns occurred either regularly or with a temporal jitter within the sequences, and participants’ attention was directed either towards or away from the auditory stimulation. Across both regular and jittered sequences during both attention and in-attention, pattern repetitions led to increased sustained activity throughout the sequence, evoked a characteristic positivity-negativity complex in the event-related potential, and enhanced inter-trial phase coherence of low-frequency oscillatory activity time-locked to repeating pattern onsets. While regularity only had a minor (if any) influence, attention significantly strengthened pattern repetition perception, which was consistently reflected in all three EEG markers. These findings suggest that the detection of pattern repetitions within continuous sounds relies on a flexible mechanism that is robust against in-attention and temporal irregularity, both of which typically occur in naturalistic listening situations. Yet, attention to the auditory input can enhance processing of repeating patterns and improve repetition detection.

## Introduction

Detection of repeating patterns is crucial for efficient perception of sounds that continuously unfold in time (1,2). Especially in complex listening situations that involve several simultaneously active sound sources, recognition of familiar sound patterns facilitates the segregation of sound streams and enables rapid adaptive reactions to change in the environment (3–7). There is compelling evidence that the human auditory system is exceptionally sensitive to pattern repetitions in sounds, even when the acoustic signal contains only minimal spectro-temporal structure such as in the case of (periodic) white noise (8–13).

Numerous studies have investigated both behavioural and neural correlates of pattern repetition detection in continuous streams of complex and meaningless sounds, including white noise (8,13–21), sequences of tone pips (22–29), “tone clouds” (30,31), and “correlated noise” (32). Besides above-chance behavioural detection of repetitions with a performance that is comparable to an ideal observer model (22), characteristic changes in several electroencephalography (EEG)/magnetoencephalography (MEG) markers were found to reflect (automatic) repetition detection: First, compared with random stimulus sequences (without pattern repetitions), an increase in magnitude of the sustained response typically occurred relative to the first pattern repetition within a sequence (22,24,25,27–29). Second, repeating pattern onsets (within the continuous sound) evoked a characteristic negativity in the event-related potential (ERP; 15,16,18,20,26,33), in some studies preceded by an early positivity (20,26,33). Finally, pattern repetitions within a sound sequence enhanced inter-trial phase coherence (ITPC) of low-frequency neural oscillations (relative to sequences without repetitions; 15,16,19,20,25). While in most studies ITPC may have at least partly reflected a sequence of ERPs evoked by periodically repeating pattern onsets, there is also evidence for synchronisation of oscillatory activity beyond the frequency of (isochronous) pattern occurrence in the stimulation (19).

A growing number of studies has moved beyond using strictly isochronous pattern repetitions and asking participants to complete an active repetition detection task. In fact, any mechanism that can possibly support pattern repetition detection in real-life listening situations should be somewhat tolerant to listeners’ in-attention and temporal irregularity with regard to pattern occurrence in the stimulus stream. Several studies showed that this is indeed the case: A negativity in the ERP was elicited relative to the onset a repeating pattern in white noise not only when participants’ attention was focussed on the auditory stimuli, but also when they were presented with the noise sequences while reading a book (33), performing a visual distractor task (15), and even during sleep (16). Similarly, pattern repetitions in white noise and sequences of tone pips led to an increase in ITPC while participants were asleep (16) or focussed on a concurrent visual task (25). A repetition-related increase in sustained response magnitude to sequences of tone pips in the absence of listeners’ attention to the auditory stimulation was reported by some studies (22,27– 29), but not by others (25). Only one study investigated the role of temporal regularity for the detection of pattern repetitions in tone pip sequences: Hodapp & Grimm (2021) found that a negativity time-locked to repeating pattern onsets was elicited consistently across temporally regular and jittered sequences, whereas the earlier positivity occurred only in regular sequences. They therefore argued that, while the negative component is related to the repetition of a specific pattern (irrespective of temporal regularity), the additional positive component in regular, temporally predictable sequences reflects neural entrainment to the periodic stimulus rhythm and anticipation of upcoming pattern occurrences (26).

Taken together, neither attention nor temporal regularity appears to be indispensable for the successful detection of repeating patterns in continuous sounds. However, since earlier studies only focussed on either of the two factors and not always directly compared different levels of attention or regularity, less is known about the interaction between attention and regularity and about whether they substantially modulate repetition perception. For instance, it remains unclear whether irregular repetitions could also be detected in the absence of attention, and whether attention and regularity improve (or in-attention and irregularity impair) the detection of pattern repetitions. Moreover, previous findings revealed some discrepancy with regard to the influence of attention on different repetition-related EEG markers (often analysed only in separate studies). One study analysed both sustained activity and ITPC within the same dataset and found that temporal regularity of a sound led to an increase of ITPC irrespective of the listeners’ attentional state, while an increase in sustained activity was only observed during attention (but not during in-attention; Herrmann & Johnsrude, 2018). Therefore, the authors argued that the two markers might reflect functionally dissociable stages of repetition perception (25). The current study aims to systematically assess in a two-by-two design how attention and temporal regularity (as well as their interaction) shape pattern repetition perception and influence its different neural signatures (within the same dataset). To this end, we presented listeners with sequences of correlated noise that contained (or did not contain) repetitions of a certain sound segment, with repetitions occurring either in a temporally regular or jittered manner, while attention was directed either towards or away from the auditory stimulation. We analysed three different EEG markers that were previously related to successful repetition detection: global field power (GFP) as a measure of sustained activity throughout the sequence, ERPs time-locked to repeating pattern onsets, and ITPC. That way, we might be able to reconcile previous, partly discrepant, findings on the role of attention and regularity and provide a more comprehensive view on auditory repetition perception and its neural correlates.

## Materials and Methods

The present study is a reanalysis of a dataset that was previously used to explore a different research question, namely the formation of memories for recurring sound patterns *across* trials (34). Conversely, the current analysis investigates the perception of pattern repetitions *within* sounds.

### Participants

29 participants (26 female, three male), aged 18 to 32 years (*M* = 21.38 years, *SD* = 3.21 years), took part in the study. None of them reported impaired hearing or a history of any neurological or psychiatric disorder, and all of them had normal or corrected-to-normal vision. Participants were recruited at Leipzig University between April and July 2022. All participants were naïve regarding the purpose of the study, gave written informed consent before the start of the experiment, and received course credits for their participation. Consent forms were stored separately from the experimental data, and any personal data were pseudonymised, such that after data collection individual participants could not be identified. We obtained written approval by a local ethics committee (Ethics Advisory Board at Leipzig University; reference number: 2022.01.26_eb_128) prior to the study, and all experimental procedures were in accordance with the Declaration of Helsinki.

### Stimuli

We used sequences of correlated noise as auditory stimulus material. Correlated noise was described in detail by McDermott and colleagues (2011) and refers to randomly generated white noise sequences that were transformed using a generative model to match statistical properties of natural sounds. Stimulus sequences were created using the Gaussian Sound Synthesis Toolbox (http://mcdermottlab.mit.edu/Gaussian_Sound_Code_for_Distribution_v1.1) in Matlab (version R2021a; The MathWorks Inc., USA), with a duration of 3500 ms, including 5-ms onset and offset ramps (half-Hanning windows). Transformation of the white noise sequences resulted in correlated noise sequences with a correlative structure, i.e., adjacent sampling points along the temporal and spectral dimension were correlated with regard to their spectral energy values, and the strength of this correlation decreased with increasing distance. Decay constants were the same as in the original study (−0.065 per 20 ms and −0.075 per 0.196 octaves), such that the structure of the generated stimuli matches the correlative structure of natural sounds (5).

We created sequences of random correlated noise without repetitions (“no repetition”; no-rep) and sequences that contained repetitions (“repetition”; rep). In rep sequences, a certain 200-ms segment occurred in total six times throughout the sequence. Rep sequences were created by inserting a separately generated 200-ms sound pattern into the 3500-ms sequence. For half of the rep sequences within an experimental block the same repeating 200-ms pattern was used, whereas for the other half a new pattern was created for each sequence. As this procedure resulted in local disruptions in the correlative structure of the sound at pattern boundaries, we controlled for these local changes by inserting six (different) 200-ms segments into no-rep sequences. Cross-fading (using 5-ms half-Hanning windows centred 2.5 ms relative to the beginning and −2.5 ms relative to the end of an inserted 200-ms patterns) was used to avoid audible artefacts due to abrupt changes in the spectrum at segment boundaries. In all sequences, the time point of the first pattern onset was selected randomly between 50 and 500 ms relative to sound onset. The following pattern onsets occurred either with a constant interval of 300 ms (regular) or variable intervals between 50 and 550 ms (jittered) between patterns. In jittered sequences, intervals between patterns were chosen randomly, with the restriction that the duration of two consecutive inter-pattern intervals must differ by at least 50 ms. Stimulus sequences are illustrated in Fig 1 (panel A), and audio examples can be found in the online supplemental material (https://osf.io/xn9t4/?view_only=582f31e68ff646afacfb0f4135f8bd83).

**Fig 1.**
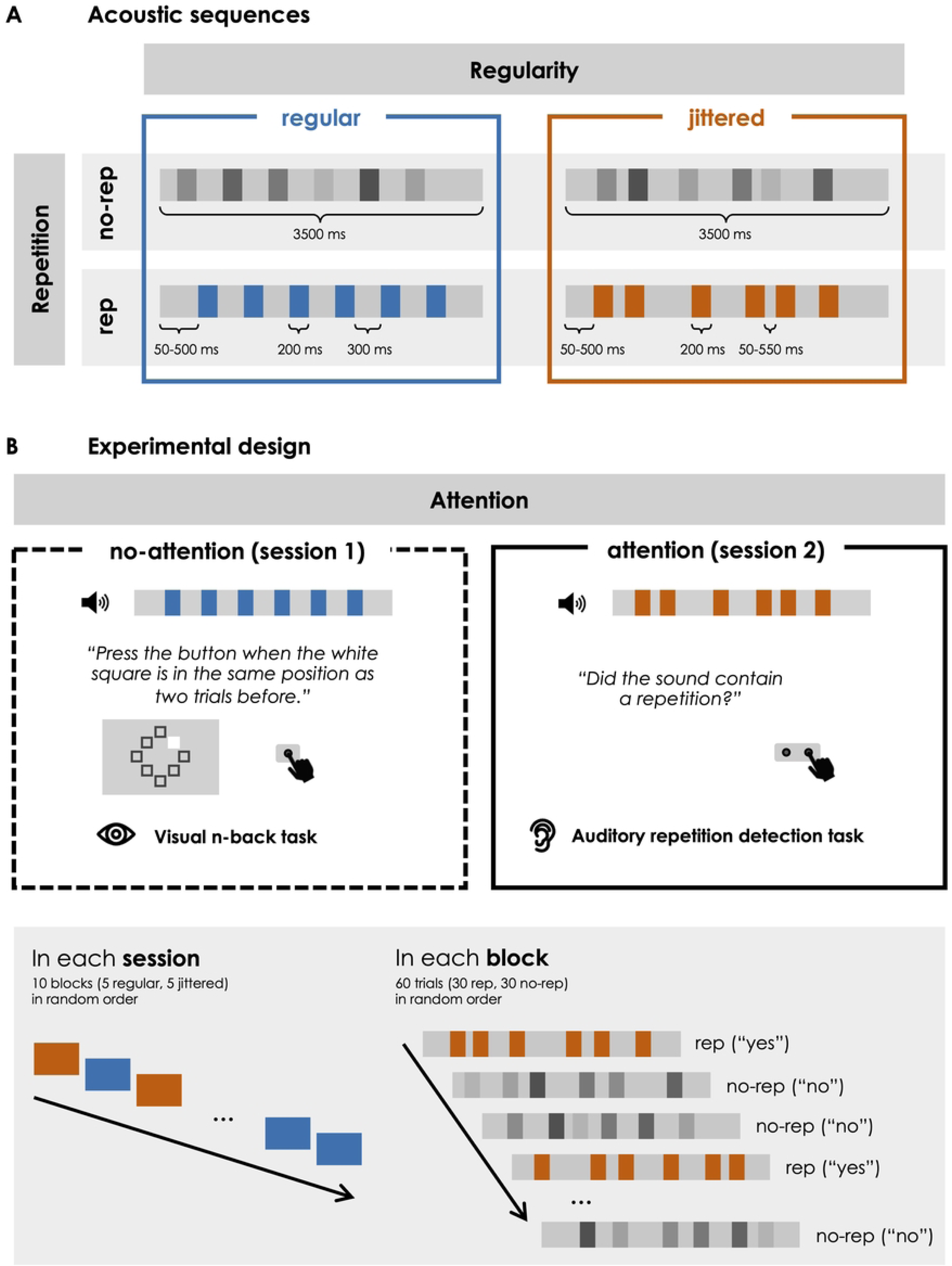
Illustration of the study design. A: Illustration of regular and jittered acoustic stimulus sequences with and without pattern repetitions. B: Experimental design. Participants took part in two EEG sessions. In the first session, their attention was directed away from the auditory stimulation, and in the second session, they were instructed to focus on repetitions in the sounds. Both sessions consisted of ten blocks in a random order, each of which contained 30 trials.

### Procedure

Participants completed two EEG sessions on separate days (with on average 13 days in between). In the first session, listeners’ attention was directed away from the auditory sequences (no-attention), which they were instructed to ignore while performing a visual distractor task that required continuous monitoring of the visual stimulation. In the second session, their attention was directed towards the auditory sequences (attention) by a repetition detection task, which required them to indicate in each trial whether the sequence contained a repetition. The fixed session order served to avoid active knowledge about the repetitions in the auditory sequences during the no-attention session after participants performed the auditory repetition detection task in the session before. In each session, they completed five blocks with regular and five blocks with jittered sequences in a random order, with breaks between blocks as required. Each block consisted of 60 randomly ordered auditory sequences, 50 % of which were rep and no-rep sequences, respectively. In 50 % of the rep sequences per block, the repeating pattern was the same across trials within the block, whereas the remaining rep sequences contained a repeating pattern that occurred in only one trial throughout the experiment. Between two consecutive sequences, silent intervals ranged between 2175 and 2625 ms in duration (in steps of 50 ms). The experimental design is illustrated in Fig 1 (panel B).

The visual display in the no-attention session consisted of eight squared dark-grey frames (width/height: 0.50° visual angle) arranged in a circle (radius: 2.11° visual angle) on a grey background at equal distance from a white fixation cross. In each of the 240 visual trials per block, a white square appeared at one of the eight frame positions for 150 ms. Participants were asked to fixate the cross in the centre of the screen and press a button a quickly as possible whenever the white square appeared at the same frame position as two trials before. The first five trials of each block were always non-targets, and 2-back targets occurred randomly in 10 % of the trials, each of which was followed by at least two non-targets. While square positions were chosen randomly for non-target trials, targets occurred equally often at each position. The visual stimulus onset asynchrony ranged between 1425 and 1575 ms (in steps of 10 ms), and visual stimulation had no temporal relationship with the auditory stimulation. Auditory stimulation began five seconds after the visual stimulation at the beginning of each block. At the beginning of the session, participants completed a short training block without concurrent auditory stimulation, during which they received feedback about the correctness of their response in each trial. During the actual experiment, feedback (hit/false alarm rates and mean reaction time) was provided only at the end of a block.

At the beginning of the attention session, the different types of auditory sequences were introduced to the participants. An example sequence (which was not used during the actual experiment) was provided for sequences with “regular repetitions” (rep, regular), “irregular repetitions” (rep, jittered) or “no repetitions” (no-rep) and could be repeated as often as listeners wanted. They were informed that repetitions occurred in 50 % of the trials and that regular and irregular sequences occurred in separate blocks. A white fixation cross on a grey background was displayed during sound presentation, followed by the response options (“repetition”/”no repetition”) during the response interval (until a response was given or a maximum of 2000 ms expired). Participants pressed either the left or the right button (counterbalanced across participants) on a response time box with their left or right index finger, respectively. Feedback (percentage of correct responses) was again provided at the end of a block.

Participants were seated inside an acoustically and electrically shielded chamber during the experiment. Task instructions and visual stimuli were presented on a computer screen located at approximately 80 cm distance from the participants’ eyes. Auditory stimuli were delivered binaurally via headphones (Sennheiser HD-25-1, Sennheiser GmbH & Co. KG, Germany). Stimulus presentation and response registration was controlled using the Psychophysics Toolbox extension (PTB-3; (35,36) in GNU Octave (version 5.2.0), and behavioural responses were recorded with a response time box (Suzhou Litong Electronic Co., China).

### EEG data acquisition

We recorded the continuous EEG from 64 Ag/AgCl electrodes mounted in an elastic cap according to the extended 10-20 system. To record horizontal and vertical eye movements, additional electrodes were placed on the outer canthus of both eyes and above and below the right eye. Signals were also recorded from the left (M1) and right (M2) mastoid and from and electrode placed on the tip of the nose, which served for offline referencing. Offsets of all electrodes were kept below 30 μV. Signals were referenced to the CMS-DRL ground, amplified with a BioSemi ActiveTwo amplifier (BioSemi B.V., Amsterdam, The Netherlands), and digitised with a sampling rate of 512 Hz.

### Data analysis and statistical inference

Since the focus of the current study was the perception of pattern repetitions *within* a sound (and not the effect of pattern recurrence *across* trials as in the previous study; 34), all sequences with pattern repetitions were collapsed into the same condition (rep) for the present analysis. To make sure that the repetition of patterns across sequences did not bias the current results, the analysis was repeated analogously excluding sequences that contained repetitions of patterns that reoccurred across trials. This approach yielded a virtually identical pattern of results, thus we decided to include all sequences for the sake of statistical power.

### Behavioural data

Analysis of behavioural data was done in RStudio (version 4.0.2, RStudio Inc., USA). Performance in the repetition detection task in the attention session was analysed within the framework of signal detection theory (37). Trials were classified as hits when participants correctly indicated that a rep sequence contained repetitions and as false alarms when they erroneously indicated that a no-rep sequence contained repetitions. We then computed the d’ sensitivity index from hit and false alarm rates separately for regular and jittered blocks, applying a log-linear transformation (38) to avoid infinite values. To statistically test whether there was a difference in repetition detection performance between regular and jittered blocks, we compared d’ scores using a two-sided paired *t*-test, with the standard .05 alpha criterion to define statistical significance. Bayesian tests were computed, using the package “BayesFactor” (39,40), and Bayes Factors (BF_10_) are reported in addition to the frequentist statistics. BF_10_ > 3 (10) was considered moderate (strong) evidence for the alternative hypothesis and BF_10_ < 0.33 (0.1) was considered moderate (strong) evidence for the null hypothesis, in accordance with widely used conventions (41), and values in between were considered inconclusive.

### EEG data

Offline processing of EEG data was done in Matlab (version R2022b), using the EEGLAB (version 14.1.2; 42) and FieldTrip (43) toolboxes, and statistical analysis in RStudio (version 4.0.2).

#### Pre-processing

Pre-processing of EEG data was done separately for each of the two sessions per participant. After re-referencing the data to the channel on the tip of the nose, noisy channels were identified if their signal variance exceeded an absolute z-score of 3.0. These channels were excluded from pre-processing and later spherically spline interpolated. The remaining data where then high-pass and low-pass filtered using Kaiser-windowed sinc finite impulse response (FIR) filters. The cut-off for the low-pass filter was 35 Hz (transition bandwidth: 5 Hz, maximum passband deviation: 0.001, filter order: 372), while high-pass filters with different cut-offs were applied for the three EEG markers that we analysed (see below). After filtering, the continuous data were epoched from −100 to 4000 relative to sequence onset. To remove physiological and technical artefacts, an independent component analysis (ICA) was used, computed on a copy of the data filtered with a 1-Hz high-pass filter (transition bandwidth: 0.5 Hz, maximum passband deviation: 0.001, filter order: 3710) to improve signal-to-noise ratio for the decomposition. Before ICA decomposition, epochs with a peak-to-peak difference exceeding 750 µV were discarded and data were down-sampled to 128 Hz. ICA weights, obtained with an infomax algorithm implemented in the runica function in EEGLAB, were transferred to the dataset with the final pre-processing parameters. Classification of independent components was done automatically using the IC Label plugin (44), and all components classified as eye blinks, muscle or cardiac activity, line or channel noise were removed. Any auditory event within 500 ms before and after a button press or within 500 ms after a visual target in the no-attention session was excluded from the analysis to minimise the influence of motor and visual activity on auditory EEG responses.

#### Sustained response: global field power (GFP)

For the analysis of sustained activity, data were high-pass filtered (during pre-processing) with a low cut-off at 0.1 Hz (transition bandwidth: 0.2 Hz, maximum passband deviation: 0.001, filter order: 9274) to avoid filtering out slow potential shifts. From the pre-processed data, we extracted epochs that ranged from −100 to 3000 ms relative to the onset of the first pattern per sequence and baseline-corrected them to the 100-ms interval prior to first pattern onset. Epochs were discarded if their peak-to-peak difference exceeded 300 µV, and the remaining epochs were re-referenced to the average of all channels. For each participant, averages were computed for rep and no-rep sequences in each of the four attention and regularity conditions. GFP at each sampling point was computed from these within-participant averages as the root mean square (RMS) of the signal across all scalp electrodes (45).

For statistical evaluation, mean GFP was extracted for each attention and regularity condition from a time window that ranged from 500 to 3000 ms relative to the first pattern onset, i.e., from the first pattern repetition to the end of the sequence. We used a three-way repeated-measures ANOVA (implemented in the R package “ez”) with the factors Repetition (rep, no-rep), Attention (attention, no-attention), and Regularity (regular, jittered) to test whether GFP differed between sequences with and without sequences, and whether this effect is modulated by attention and regularity. Greenhouse-Geisser correction was applied whenever Mauchly’s test indicated non-sphericity (*p* < .05). A corresponding Bayesian ANOVA (46) was again computed in addition to the frequentist ANOVA. Reported BF_10_’s reflect the evidence for models that include the respective (main or interaction) effect relative to reduced matched models without the respective effect (in line with recent recommendations; 47). A significant main effect of Repetition would indicate that the brain successfully picked up the pattern repetitions within sound sequences, and a significant interaction of Repetition with Attention or Regularity would indicate that the repetition effect is modulated by the respective factor. To further elucidate the nature of the modulation by Attention or Regularity, significant (*p* < .05) two-way interactions with Repetition were followed up using (both frequentist and Bayesian) paired *t*-tests. Specifically, we computed the rep vs. no-rep contrast separately for the two levels of the modulating factor (Attention or Regularity), and subsequently compared the rep-minus-no-rep difference between the two levels (i.e., attention vs. no-attention, or regular vs. jittered).

#### Event-related potential (ERP) responses to repeating pattern onsets

For the ERP analysis, data were filtered with a 1-Hz high-pass filter (transition bandwidth: 0.5 Hz, maximum passband deviation: 0.001, filter order: 3710) in order to filter out slow potentials. Extracted epochs ranged from −100 to 500 ms relative to single pattern onsets, averaged across the second to the sixth pattern occurrence per sequence. Epochs were discarded if their peak-to-peak difference exceeded 300 µV, and no baseline correction was applied. After re-referencing to the algebraic mean of both mastoids, we computed first within-participant averages and then grand averages across participants for rep and no-rep sequences in each of the four attention and regularity conditions.

A non-parametric cluster-based permutation approach was used to determine time windows of interest for the statistical evaluation of mean ERP amplitudes. To identify clusters of significant differences in amplitude between rep and no-rep sequences at adjacent sampling points along both temporal and spatial dimension, we computed a cluster-based permutation test on rep vs. no-rep averages across the four attention and regularity conditions (48,49). Averaging across attention and regularity conditions before computing the cluster-based permutation test served to reduce the risk of biased analysis parameter choices (50). Both alpha level and cluster alpha were set to 0.05, and cluster-level significance probability was estimated using a Monte Carlo approximation with 1000 permutations. In the time range from 0 to 500 ms relative to pattern onset, we identified two time windows of interest, the first one ranging from 0 to 160 ms and corresponding to an early positivity, and the second one ranging from 190 to 380 ms and corresponding to a subsequent negativity.

Mean amplitudes were extracted from these two time windows at a fronto-central cluster of nine electrodes (F1, F2, Fz, FC1, FC2, FCz, C1, C2, Cz). Statistical evaluation was done separately for the positivity and negativity, and followed the same procedures as described above for the sustained response.

#### Inter-trial phase coherence (ITPC)

For the analysis of ITPC, data were high-pass filtered with a cut-off at 0.5 Hz (transition bandwidth: 0.5 Hz, maximum passband deviation: 0.001, filter order: 3710). Pre-processed data were epoched from −200 to 800 ms relative to single pattern onsets at the second to the sixth pattern occurrence per sequence. Epochs were demeaned, and any epoch with a peak-to-peak difference that exceeded 150 µV was discarded. Signals were averaged within the same fronto-central electrode cluster as for the ERP analysis (see above), and 1500-ms zero-padding was applied at both ends of each epoch. We then used a convolution with Morlet wavelets to extract phase information from single epochs over a frequency range from 1 to 10 Hz (in steps of 0.2 Hz), with parameters of the wavelet linearly adjusted from three to seven wavelet cycles. ITPC between epochs was computed for each participant from the results of the wavelet convolution at each sampling point in the time-frequency space, separately for rep and no-rep sequences in each of the four attention and regularity conditions. We again used a cluster-based permutation approach to determine the time-frequency window of interest for statistical evaluation. After averaging across the four attention and regularity conditions, we computed a cluster-based permutation test (rep vs. no-rep), with an alpha level and cluster alpha of 0.001 (and again using a Monte Carlo approximation with 1000 permutations to estimate cluster-level significance probability). The test revealed a broad significant cluster that ranged from 0 to 500 ms relative to pattern onset and spanned a frequency range from 1 to 4 Hz.

We extracted mean ITPC from this time-frequency window of interest for subsequent statistical evaluation, which followed the same procedures as for the analysis of sustained response and ERPs to repeating pattern onsets.

## Results

### Behavioural data

Participants detected pattern repetitions in the acoustic sequences on average above chance in both regular (*M* ± *SD* of d’: 2.01 ± 0.97) and jittered (*M* ± *SD* of d’: 1.84 ± 1.11) blocks. There was no significant difference between the two (*t*(28) = 1.92; *p* = .065; *d* = 0.36; BF_10_ = 0.99), however Bayesian evidence was inconclusive. Thus, there might in fact be a trend towards better repetition detection performance in regular than in jittered sequences, though the effect of temporal regularity seems to be rather small.

### EEG data

#### Sustained response: GFP

As displayed in Fig 2, GFP overall increased rather sharply at the beginning of a sequence before reaching a relatively sustained plateau phase throughout the rest of the sequence from around 500 ms after the first pattern onset. In regular rep sequences (across both attention conditions), we observed an additional periodic modulation of the potential during the sustained phase at the frequency of the isochronous repeating pattern onsets (i.e., 2 Hz). Any such response relative to repeating pattern onsets would be levelled out due to the random shift of pattern onsets in jittered sequences. Crucially, GFP was significantly higher in rep compared to no-rep sequences (main effect of Repetition: *F*(1, 28) = 48.39, *p* < .001, partial η^2^ = .63, BF_10_ = 2.21*10^5^), suggesting that the brain automatically picked up pattern repetitions in rep sequences. This repetition effect was modulated by attention (Repetition x Attention interaction: *F*(1, 28) = 6.80, *p* = .014, partial η^2^ = .20, BF_10_ = 0.87): While there was a significant increase in GFP for rep compared to no-rep sequences during both attention (*t*(28) = 6.65; *p* < .001; *d* = 1.24; BF_10_ = 5.05*10^4^) and in-attention (*t*(28) = 5.55; *p* < .001; *d* = 1.03; BF_10_ = 3.30*10^3^), the effect was larger when listeners’ attention was focussed on the sounds (*t*(28) = 2.61; *p* = .014; *d* = 0.48; BF_10_ = 3.34). Conversely, the influence of regularity on the repetition effect was less clear (Repetition x Regularity interaction: *F*(1, 28) = 5.41, *p* = .027, partial η^2^ = .16, BF_10_ = 0.47): The repetition effect was significant in both regular (*t*(28) = 4.67; *p* <.001; *d* = 0.87; BF_10_ = 366.93) and jittered (*t*(28) = 6.81; *p* < .001; *d* = 1.26; BF_10_ = 7.41*10^4^) blocks, and there was a trend towards a larger effect in jittered blocks, although only with inconclusive Bayesian evidence (*t*(28) = 2.33; *p* = .027; *d* = 0.43; BF_10_ = 1.96).

**Fig 2.**
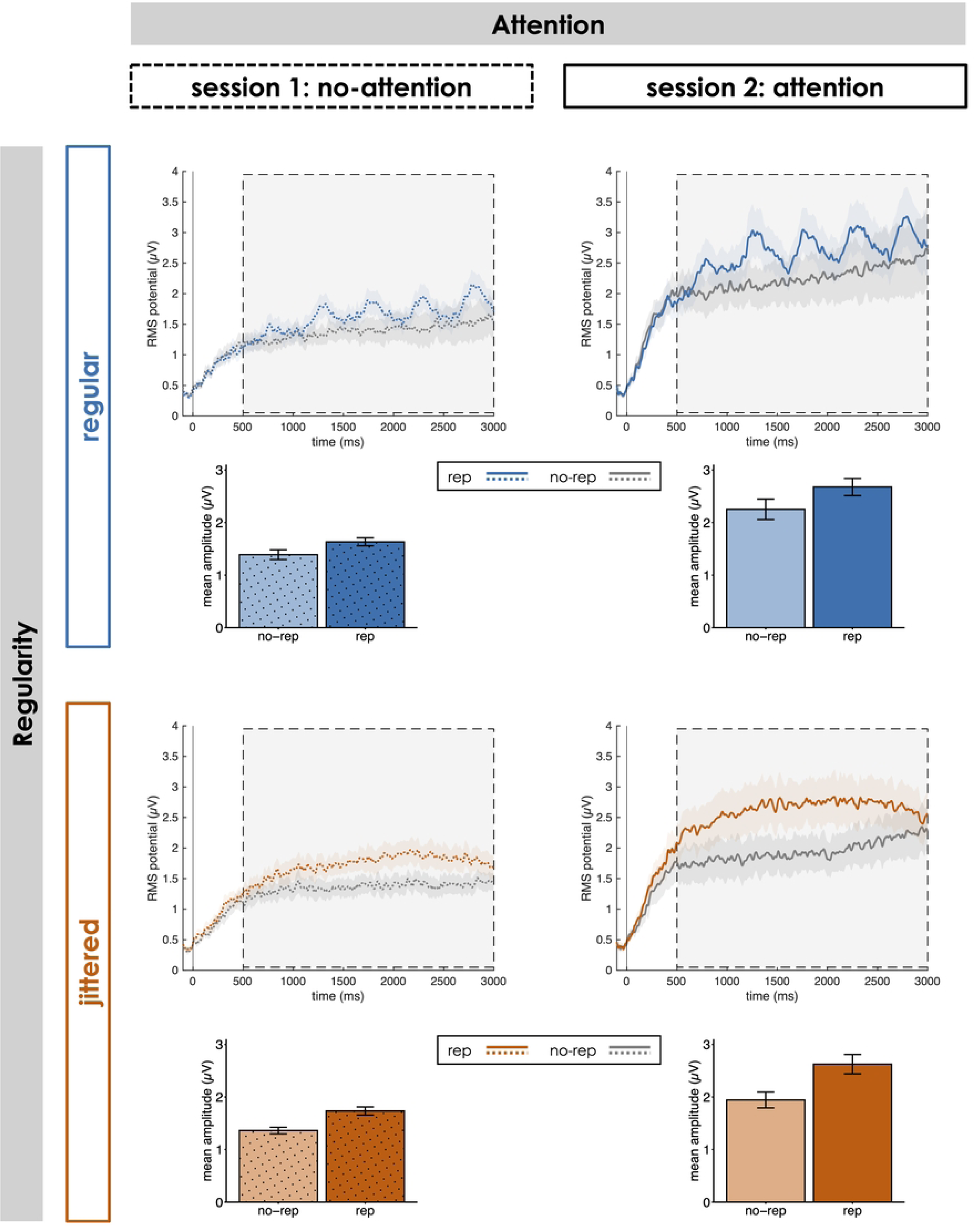
Sustained activity throughout the sequence. Global field power (GFP) relative to the onset of the first pattern occurrence per sequence (0 ms) for rep and no-rep sequences in each of the four Attention x Regularity conditions. Bar plots display mean amplitudes in the time window of interest (500 to 3000 ms; marked by the light-grey box). Shaded areas and error bars indicate ± 1 SEM.

### ERPs to repeating pattern onsets

ERPs to the onsets of the repeating pattern within a sequence are shown in Fig 3. Repeating pattern onsets in rep sequences elicited an early positivity, followed by a later negativity from around 200 ms relative to pattern onset, both with a fronto-central topography, whereas no such ERP modulation occurred for no-rep sequences. This pattern-related positivity-negativity complex was elicited consistently across all attention and regularity conditions, with differences only in the latency of the positivity: While the onset of the positivity was around pattern onset in jittered sequences, it was shifted forwards in regular sequences, likely related to anticipation of upcoming pattern repetitions in temporally regular and predictable sequences. For both positivity (0-160 ms) and negativity (190-380 ms) effects of Repetition, Attention and Regularity pointed into the same directions: Mean amplitudes were larger (i.e., more positive or negative, respectively) in rep than in no-rep sequences (main effect of Repetition: positivity: *F*(1, 28) = 123.29, *p* < .001, partial η^2^ = .81, BF_10_ = 8.30*10^47^; negativity: *F*(1, 28) = 182.74, *p* < .001, partial η^2^ = .87, BF_10_ = 2.91*10^55^). While the repetition effect was not significantly modulated by regularity (Repetition x Regularity interaction: positivity: *F*(1, 28) = 0.21, *p* = .654, partial η^2^ = .01, BF_10_ = 0.32; negativity: *F*(1, 28) = 1.57, *p* = .220, partial η^2^ = .05, BF_10_ = 0.27), there was a substantial influence of attention (Repetition x Attention interaction: positivity: *F*(1, 28) = 25.99, *p* < .001, partial η^2^ = .48, BF_10_ = 1.49*10^3^; negativity: *F*(1, 28) = 52.98, *p* < .001, partial η^2^ = .65, BF_10_ = 4.46*10^7^): Amplitudes differed significantly between rep and no-rep sequences during both attention (positivity: *t*(28) = 12.57; *p* < .001; *d* = 2.33; BF_10_ = 1.54*10^9^; negativity: *t*(28) = 13.88; *p* < .001; *d* = 2.58; BF_10_ = 1.53*10^11^) and in-attention (positivity: *t*(28) = 7.71; *p* < .001; *d* = 1.43; BF_10_ = 6.39*10^5^; negativity: *t*(28) = 9.70; *p* < .001; *d* = 1.80; BF_10_ = 5.48*10^7^), but an attentional focus on the auditory sequences increased this repetition effect (positivity: *t*(28) = 5.10; *p* < .001; *d* = 0.95; BF_10_ = 1.07*10^3^; negativity: *t*(28) = 7.28; *p* < .001; *d* = 1.35; BF_10_ = 2.30*10^5^).

**Fig 3.**
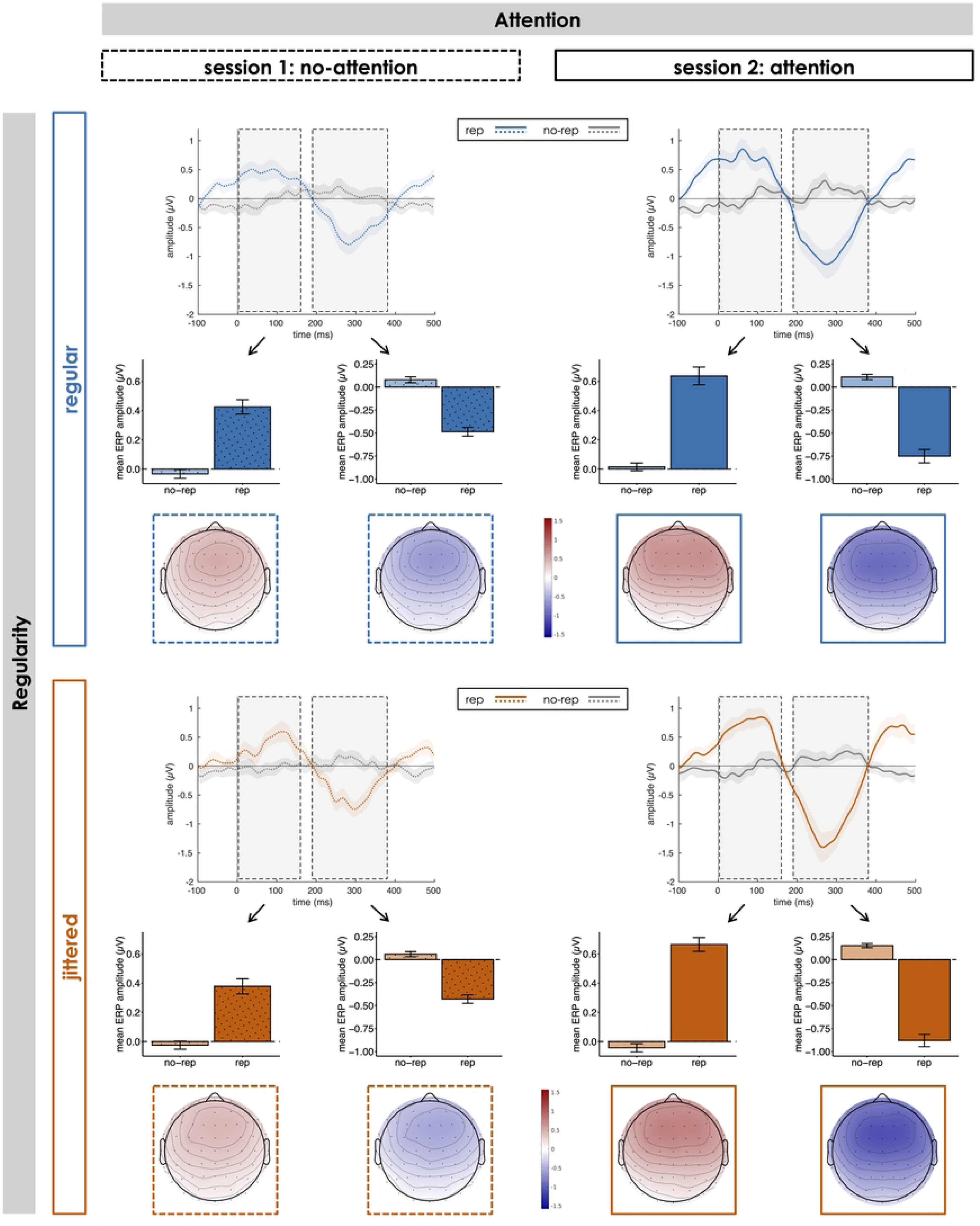
Event-related responses to repeating pattern onsets. Event-related potentials (ERPs) relative to the onset of repeating patterns at position 2 to 6 within the sequence (0 ms) for rep and no-rep sequences in each of the four Attention x Regularity conditions. Bar plots display mean amplitudes in the time windows of interest (early positivity: 0 to 160 ms; later negativity: 190 to 380 ms; marked by light-grey boxes) at a fronto-central electrode cluster. Topographies show the rep-minus-no-rep difference potential in the respective time window of interest. Shaded areas and error bars indicate ± 1 SEM.

### ITPC

As shown in Fig 4, pattern repetitions within a sound sequence led to an increase in ITPC of neural oscillations (compared to no-rep sequences). ITPC was strongest around the onsets of the repeating pattern for low frequencies around the rates of pattern occurrence in the stimulation. In regular sequences, the ITPC peak appeared more focal along the frequency dimension, which reflects the strict 2-Hz periodicity in the stimulation compared to jittered sequences that comprise a broader range of frequencies (1.33 to 4 Hz). Statistical evaluation of mean ITPC between 0 and 500 ms relative to pattern onset showed that phase coherence of 1-4 Hz oscillations was overall stronger in rep than in no-rep sequences (main effect of Repetition: *F*(1, 28) = 69.61, *p* < .001, partial η^2^ = .71, BF_10_ = 8.91*10^23^). The repetition effect was significantly modulated by attention (Repetition x Attention interaction: *F*(1, 28) = 63.35, *p* < .001, partial η^2^ = .69, BF_10_ = 5.12*10^9^): The increase in ITPC for rep compared to no-rep sequences was significant during both attention (*t*(28) = 9.25; *p* <.001; *d* = 1.72; BF_10_ = 2.09*10^7^) and in-attention (*t*(28) = 4.37; *p* < .001; *d* = 0.81; BF_10_ = 178.13), but the difference was substantially larger during attention (*t*(28) = 7.96; *p* < .001; *d* = 1.48; BF_10_ = 1.14*10^6^). Similarly, regularity also influenced the magnitude of the repetition effect (Repetition x Regularity interaction: *F*(1, 28) = 7.03, *p* = .013, partial η^2^ = .20, BF_10_ = 2.09): While there was a significant repetition effect in both regular (*t*(28) = 7.57; *p* < .001; *d* = 1.41; BF_10_ = 4.56*10^5^) and jittered (*t*(28) = 7.55; *p* < .001; *d* = 1.40; BF_10_ = 4.35*10^5^) blocks, the effect was larger in regular blocks (*t*(28) = 2.65; *p* = .013; *d* = 0.49; BF_10_ = 3.65).

**Fig 4.**
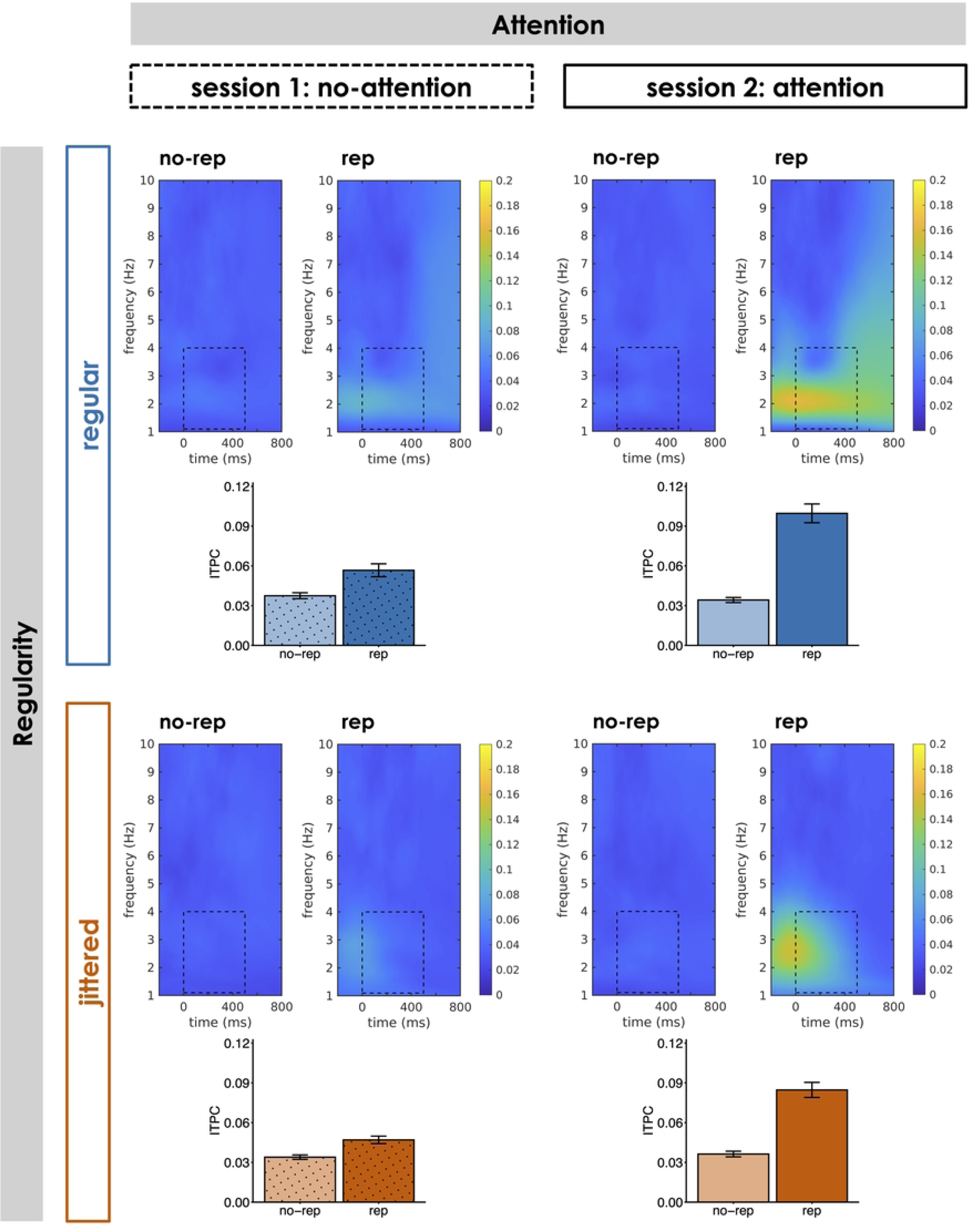
Phase coherence of neural oscillations. Inter-trial phase coherence (ITPC) over frequencies and time relative to the onset of repeating patterns at position 2 to 6 within the sequence (0 ms) at a fronto-central electrode cluster for rep and no-rep sequences in each of the four Attention x Regularity conditions. Bar plots display mean ITPC between 1 and 4 Hz in a time window from 0 to 500 ms relative to pattern onset (marked by dotted lines). Error bars indicate ± 1 SEM.

## Discussion

The current study set out to test whether and how listeners’ attention and the temporal regularity of pattern occurrence within a continuous sound sequence modulate pattern repetition perception. We presented listeners with sequences of correlated noise that contained or did not contain repetitions of a certain sound segment. Pattern repetitions within a sequence were either temporally regular or jittered, and listeners’ attention was either directed towards or away from the sounds during stimulus presentation. Besides behavioural repetition detection (when participants attended to the sounds), we measured repetition perception and its modulation by attention and regularity by means of three different EEG signatures: sustained activity throughout the full sequence (from repetition onset), ERPs and ITPC time-locked to repeating pattern onsets.

Overall, listeners were able to behaviourally detect repetitions well above chance (when they attended to the sounds), and successful repetition detection was reflected consistently in all three neural markers across attention and regularity conditions. Concretely, repetitions of a specific pattern within a continuous acoustic stimulus led to an increase in sustained activity from the first pattern repetition through the end of the sequence (for consistent previous results, see: 22,24,25,27–29), a characteristic positivity-negativity complex in the ERP time-locked to repeating pattern onsets (15,16,18,20,26,33), and enhanced ITPC of low-frequency (1-4 Hz) oscillations (15,16,19,20,25). Notably, besides replicating findings of different earlier studies all within the same dataset, we could demonstrate automatic detection of irregular, unpredictable pattern repetitions while listeners focussed on a demanding visual distractor task. Thus, we show that not only strict periodicities (15,16,22,25,27–29,33), but also more irregular pattern repetitions within continuous auditory input are processed pre-attentively. This suggests that repetition detection does not rely on a merely temporal mechanism (i.e., the detection of an autocorrelation with a fixed time lag in the acoustic signal), but on a continuous comparison between the current input and a sensory representation of a recently presented sound segment.

While pattern repetitions were detected automatically in both regular and jittered sequences during both attention and in-attention, repetition perception was substantially modulated by both factors. Our two-by-two within-subject design allowed to directly compare different levels of attention and regularity, and to show that an attentional focus onto the sounds substantially enhanced repetition perception. The repetition effect (i.e., the difference between sequences with and without pattern repetitions) was larger during attention than during in-attention to the auditory stimulation across all three neural markers. In contrast, earlier studies had suggested rather comparable magnitudes of the repetition effect between attention and no-attention as reflected in sustained activity (27), ERPs (15,16,33), and ITPC (25). However, most of these studies did either not compare attention conditions directly (15,16,22), used a between-subject design (27), or controlled attention less strictly (33). We argue that attention to the stimulus sequences (and, in particular, potential repetitions therein) enhances perceptual representation of the sound and thereby facilitates repetition detection. Sharpened short-term representations of the repeating pattern through attention may in turn boost robust memory formation for specific patterns that recur across multiple trials at a longer time scale (and potentially higher level of abstraction), which has been demonstrated previously (8,13–17,21,23,24,30–32).

Conversely, the influence of temporal regularity on repetition perception appeared somewhat less clear and consistent across different neural markers: While there was no difference in amplitude and morphology of the ERP to repeating pattern onsets between regular and jittered sequences, the repetition effect tended to be smaller for regular sequences in terms of sustained activity, but larger in terms of ITPC. The absence of a regularity-related difference in the ERP effect is only partly in agreement with the results of a previous study by Hodapp & Grimm (2021), who reported no difference in amplitude of the negative ERP component between regular and jittered pattern repetitions, whereas the early positive ERP component exclusively occurred in the regular condition. By contrast, the occurrence of both components across regular and jittered sequences in our data suggests that positivity and negativity do not subserve different functions (e.g., tracking of stimulus periodicity vs. detection of repeating pattern onsets), but rather that the positivity-negativity complex as a whole is related to pattern repetition detection. Nevertheless, the (descriptive) forward shift of the onset of the positivity for regular (compared to jittered) pattern onsets may indicate that anticipation of upcoming pattern occurrences in predictable sequences is reflected in the latency (but not in the magnitude) of the ERP response. If anticipation of upcoming pattern onsets indeed modulates the time course of the ERP such that the early positivity reaches into a time window before actual pattern onset, baseline correction could introduce amplitude differences between regular and irregular sequences by differentially shifting the whole positivity-negativity complex into a negative or positive direction (which may also explain discrepancies with regard to the presence of the early positivity in earlier studies, e.g., 15,26). A similar interpretation may hold for the stronger ITPC effect we observed for regular than for jittered sequences: The strict periodicity in the stimulation allowed for (additional) entrainment of brain responses to the stimulus rhythm and for temporal prediction of the next pattern onset, which was not possible in unpredictable jittered sequences. Importantly, the presence of a significant repetition-related ITPC increase for jittered sequences suggests that the phase alignment of EEG responses cannot be explained merely by entrainment to the stimulus periodicity. Instead, synchronisation of neural responses relative to repeating pattern onsets occurred irrespective of their temporal regularity, possibly achieved via phase-reset of ongoing oscillations (19,51). Finally, there was a trend towards a larger repetition effect in sustained activity for jittered compared to regular sequences, which may seem counterintuitive at first glance. Especially in the attention condition, this effect seems to be driven by a GFP difference between regular and jittered sequences without pattern repetitions, whereas mean GFP was (descriptively) fairly similar for sequences with repetition. This suggests that there might have been rudimentary processing of local disruptions in the correlative structure of the stimulus sequences when they occurred strictly periodically (but not when their occurrence was jittered and unpredictable), which in turn decreased the difference between rep and no-rep sequences (i.e., the repetition effect).

Unlike Herrmann & Johnsrude (2018), we did not find evidence for a distinct pattern of attention modulation between sustained activity and phase coherence of neural oscillations. If anything, our data provide more evidence for an attention modulation of the repetition effect in ITPC than in GFP, whereas Herrmann & Johnsrude (2018) reported an attention effect only for sustained activity, but not for ITPC (i.e., neural synchronisation). They proposed that the distinct susceptibility of sustained activity and neural synchronisation to the influence of attention may indicate that the two neural markers reflect dissociable processes, such that neural synchronisation is related to an early attention-independent sensory process and sustained activity to a more abstract representation of structure in sounds that requires attention (25). While this does not preclude that different EEG markers reflect functionally nuanced processes that contribute to (automatic) repetition perception, our data suggest that all of them underlie a similar modulatory influence by attention. Different weighting of putative subprocesses and their susceptibility to attention (and possibly regularity) modulation might rather arouse from subtle differences in the experimentally created listening context (e.g., specific stimulus material and distractor task).

In summary, our study replicates the results of earlier studies that showed rapid and automatic detection of pattern repetitions within continuous acoustic sequences. Crucially, pattern repetitions are processed pre-attentively even if there is no temporal regularity that could act as a cue for upcoming (predictable) pattern occurrences. This suggests that repetition perception relies on a mechanism that flexibly adapts to varying contextual demands, such as they occur in naturalistic listening situations. Yet, an attentional focus on the auditory input enhances sensory representation of repeating patterns and facilitates repetition detection.

## Acknowledgements

The authors thank Benjamin Eichenberger for help with EEG data collection.

## Supporting information

Data of individual participants and scripts to reproduce the statistical analysis reported in the manuscript can be found here: https://osf.io/xn9t4/?view_only=582f31e68ff646afacfb0f4135f8bd83 Further data and materials are available from the corresponding author upon reasonable request.

## Notes

### Competing Interest Statement

The authors have declared no competing interest.

## References

1. Chait M. How the brain discovers structure in sound sequences. Acoust Sci Technol. 2020;41(1): 48–53.

2. Maravall M, Ostojic S, Pressnitzer D, Chait M. More than the Sum of its Parts: Perception and Neuronal Underpinnings of Sequence Processing. Neuroscience. 2018;389: 1–3.

3. Bendixen A. Predictability effects in auditory scene analysis: a review. Front Neurosci. 2014;8(8): 1–16.

4. Masutomi K, Barascud N, Kashino M, McDermott JH, Chait M. Sound segregation via embedded repetition is robust to inattention. J Exp Psychol Hum Percept Perform. 2016;42(3): 386–400.

5. McDermott JH, Wrobleski D, Oxenham AJ. Recovering sound sources from embedded repetition. Proc Natl Acad Sci. 2011;108(3): 1188–93.

6. Winkler I, Denham SL, Nelken I. Modeling the auditory scene: predictive regularity representations and perceptual objects. Trends Cogn Sci. 2009;13(12): 532–40.

7. Woods KJP, McDermott JH. Schema learning for the cocktail party problem. Proc Natl Acad Sci. 2018;115(14): E3313–3322.

8. Agus TR, Thorpe SJ, Pressnitzer D. Rapid Formation of Robust Auditory Memories: Insights from Noise. Neuron. 2010;66(4): 610–618.

9. Guttman N, Julesz B. Lower Limits of Auditory Periodicity Analysis. J Acoust Soc Am. 1963;35(4): 610–610.

10. Kaernbach C. On the consistency of tapping to repeated noise. J Acoust Soc Am. 1992;92(2): 788–93.

11. Kaernbach C. Temporal and spectral basis of the features perceived in repeated noise. J Acoust Soc Am. 1993;94(1): 91–7.

12. Kaernbach C. The Memory of Noise. Exp Psychol. 2004;51(4): 240–248.

13. Viswanathan J, Rémy F, Bacon-Macé N, Thorpe SJ. Long Term Memory for Noise: Evidence of Robust Encoding of Very Short Temporal Acoustic Patterns. Front Neurosci;10: 1–11

14. Agus TR, Pressnitzer D. The detection of repetitions in noise before and after perceptual learning. J Acoust Soc Am. 2013;134(1): 464–473.

15. Andrillon T, Kouider S, Agus T, Pressnitzer D. Perceptual Learning of Acoustic Noise Generates Memory-Evoked Potentials. Curr Biol. 2015;25(21): 2823–2829.

16. Andrillon T, Pressnitzer D, Léger D, Kouider S. Formation and suppression of acoustic memories during human sleep. Nat Commun. 2017;8(1): 179.

17. Dauer T, Henry MJ, Herrmann B. Auditory perceptual learning depends on temporal regularity and certainty. J Exp Psychol Hum Percept Perform. 2022;48(7): 755–770.

18. Kaernbach C, Schröger E, Gunter TC. Human event-related brain potentials to auditory periodic noise stimuli. Neurosci Lett. 1998;242(1): 17–20.

19. Luo H, Tian X, Song K, Zhou K, Poeppel D. Neural Response Phase Tracks How Listeners Learn New Acoustic Representations. Curr Biol. 2013;23(11): 968–974.

20. Ringer H, Schröger E, Grimm S. Within- and between-subject consistency of perceptual segmentation in periodic noise: A combined behavioral tapping and EEG study. Psychophysiology. 2023;60(2): e14174.

21. Song K, Luo H. Temporal Organization of Sound Information in Auditory Memory. Front Psychol. 2017;8: 999.

22. Barascud N, Pearce MT, Griffiths TD, Friston KJ, Chait M. Brain responses in humans reveal ideal observer-like sensitivity to complex acoustic patterns. Proc Natl Acad Sci. 2016;113(5): E616– 625.

23. Bianco R, Harrison PM, Hu M, Bolger C, Picken S, Pearce MT, et al. Long-term implicit memory for sequential auditory patterns in humans. eLife. 2020;9: e56073.

24. Herrmann B, Araz K, Johnsrude IS. Sustained neural activity correlates with rapid perceptual learning of auditory patterns. NeuroImage. 2021;238: 118238.

25. Herrmann B, Johnsrude IS. Neural Signatures of the Processing of Temporal Patterns in Sound. J Neurosci. 2018;38(24): 5466–5477.

26. Hodapp A, Grimm S. Neural signatures of temporal regularity and recurring patterns in random tonal sound sequences. Eur J Neurosci. 2021;53(8): 2740–2754.

27. Sohoglu E, Chait M. Detecting and representing predictable structure during auditory scene analysis. eLife. 2016;5: e19113.

28. Southwell R, Chait M. Enhanced deviant responses in patterned relative to random sound sequences. Cortex. 2018;109: 92–103.

29. Southwell R, Baumann A, Gal C, Barascud N, Friston K, Chait M. Is predictability salient? A study of attentional capture by auditory patterns. Philos Trans R Soc B Biol Sci.;372(1714): 20160105.

30. Agus TR, Pressnitzer D. Repetition detection and rapid auditory learning for stochastic tone clouds. J Acoust Soc Am. 2021;150(3): 1735–1749.

31. Kumar S, Bonnici HM, Teki S, Agus TR, Pressnitzer D, Maguire EA, et al. Representations of specific acoustic patterns in the auditory cortex and hippocampus. Proc R Soc B Biol Sci. 2014;281(1791): 20141000.

32. Ringer H, Schröger E, Grimm S. Perceptual Learning and Recognition of Random Acoustic Patterns. Audit Percept Cogn. 2022;5(3–4): 259–281.

33. Berti S, Schröger E, Mecklinger A. Attentive and pre-attentive periodicity analysis in auditory memory: an event-related brain potential study. Neuroreport. 2000;11(9): 1883–1887.

34. Ringer H, Schröger E, Grimm S. Perceptual learning of random acoustic patterns: Impact of temporal regularity and attention. bioRxiv. 2023. doi: 10.1101/2023.03.13.532336

35. Brainard DH. The Psychophysics Toolbox. Spat Vis. 1997;10(4): 433–436.

36. Kleiner M, Brainard D, Pelli DG, Ingling A, Murray R, Broussard C. What’s new in Psychtoolbox-3? Perception 36 ECVP Abstract Supplement. 2007.

37. Macmillan NA. Signal Detection Theory. In: Smelser NJ, Baltes PB, editors. International Encyclopedia of the Social & Behavioral Sciences. Oxford: Pergamon; 2001. pp. 14075–14078.

38. Hautus MJ, Lee A. Estimating sensitivity and bias in a yes/no task. Br J Math Stat Psychol. 2006;59(2): 257–273.

39. Morey RD, Rouder JN. Bayes factor approaches for testing interval null hypotheses. Psychol Methods. 2011;16(4): 406–419.

40. Rouder JN, Speckman PL, Sun D, Morey RD, Iverson G. Bayesian t tests for accepting and rejecting the null hypothesis. Psychon Bull Rev. 2009;16(2): 225–237.

41. Lee MD, Wagenmakers EJ. Bayesian Cognitive Modeling: A Practical Course. Cambridge University Press; 2014.

42. Delorme A, Makeig S. EEGLAB: an open source toolbox for analysis of single-trial EEG dynamics including independent component analysis. J Neurosci Methods. 2004;134(1): 9–21.

43. Oostenveld R, Fries P, Maris E, Schoffelen JM. FieldTrip: open source software for advanced analysis of MEG, EEG, and invasive electrophysiological data. Comput Intell Neurosci. 2011: 1–9.

44. Pion-Tonachini L, Kreutz-Delgado K, Makeig S. ICLabel: An automated electroencephalographic independent component classifier, dataset, and website. NeuroImage. 2019;198: 181–197.

45. Murray MM, Brunet D, Michel CM. Topographic ERP Analyses: A Step-by-Step Tutorial Review. Brain Topogr. 2008;20(4): 249–264.

46. Rouder JN, Morey RD, Speckman PL, Province JM. Default Bayes factors for ANOVA designs. J Math Psychol. 2012;56(5): 356–374.

47. Bergh D van den, Doorn J van, Marsman M, Draws T, Kesteren EJ van, Derks K, et al. A tutorial on conducting and interpreting a Bayesian ANOVA in JASP. LAnnee Psychol. 2020;120(1): 73–96.

48. Maris E. Statistical testing in electrophysiological studies. Psychophysiology. 2012;49(4): 549– 565.

49. Maris E, Oostenveld R. Nonparametric statistical testing of EEG- and MEG-data. J Neurosci Methods. 2007;164(1): 177–190.

50. Luck SJ, Gaspelin N. How to get statistically significant effects in any ERP experiment (and why you shouldn’t). Psychophysiology. 2017;54(1): 146–157.

51. Calderone DJ, Lakatos P, Butler PD, Castellanos FX. Entrainment of neural oscillations as a modifiable substrate of attention. Trends Cogn Sci. 2014;18(6): 300–309.

